# Genomic Insights into Derived Dwarfism and Exudivory in a Genus (*Callithrix*) of the World’s Smallest Anthropoid Monkeys

**DOI:** 10.1101/2025.05.19.654637

**Authors:** Joanna Malukiewicz, Vanner Boere, Nelson H. A. Curi, Jorge A. Dergam, Claudia Igayara de Souza, Silvia B. Moreira, Patricia A. Nicola, Marcelo Passamani, Luiz C. M. Pereira, Ricardo C.H. del Rosario, Alcides Pissinatti, Daniel L. Silva, Ita de Oliveira e Silva, Carlos R. Ruiz-Miranda, Reed A. Cartwright, Christian Roos, Anne C. Stone

## Abstract

The primate Callitrichidae family represents the smallest anthropoid primates, which are known to possess dietary specializations for eating viscous plant exudates (exudivory), and as callitrichids *Callithrix* marmosets push these biological traits to extremes. Using low-coverage whole genome sequencing of *Callithrix* species, we investigated *Callithrix* evolutionary history, species genetic diversity, and the genomic basis of derived small body size and exudivory. Our phylogenetic species tree shows that *Callithrix* likely originated in southeastern Brazil and migrated northward to the semi-arid regions of central and northeastern Brazil. We show that there is greater genetic similarity between smaller and more exudivorous *Callithrix* species relative to larger and less exudivorous species, and that the genus likely experienced extensive past reticulation. Based on these results, we argue that the *Callithrix* migration from the Atlantic Forest biome to the relatively more extreme environments of Cerrado/Caatinga biomes progressively reduced body size and increased exudivory specialization in *Callithrix* marmosets. We also identified species-specific candidate genes under putative positive selection for traits that include body growth, bone development, reproduction, fat metabolism, insulin signaling, and liporegulation. In conclusion, we increased genomic resources for understudied *Callithrix* species, while providing the most comprehensive overview of marmoset genomic diversity to-date, and identify several candidate genes under positive selection for traits related to small body size and exudivory. Future studies should sample wild marmosets more widely and perform higher-coverage genomic sequencing to better understand how specific genomic variants may impact the evolution of key *Callithrix* biological traits.

## Introduction

As members of the Callitrichidae family, *Callithrix* marmosets, which are native to Brazil’s semi-arid and humid coastal regions (Fig. 1A), possess a suite of distinctive biological traits that are considered unique among anthropoid primates (Harris et al., 2013; Consortium, 2014; Malukiewicz et al., 2020). Specifically, the diminutive body size that characterizes callitrichids was derived from a common larger-bodied ancestor (Marroig and Cheverud, 2010), with average callitrichid body mass ranging from 110g-600g (Smith and Jungers, 1997; Malukiewicz et al., 2024). The average body mass of *Callithrix* species (250g-450g) falls on the lower end of the callitrichid range, which makes these marmosets some of the world’s smallest anthropoid primates. Many callitrichids also employ a dietary strategy of exudivory where they nutritionally exploit viscous plant exudates composed of hard to digest polysaccharides (Nash, 1986; Smith, 2010). Exudivory is frequently associated with small size in primates (Nash, 1986; Smith and Jungers, 1997). Additionally, some highly exudivorous *Callithrix* species possess specialized dietary, dental, digestive, and cranio-muscular morphological modifications (Rylands and Faria, 1993; de Souza, 2016; Malukiewicz et al., 2020, 2024). Finally, birthing of dizygotic twins defines the reproductive biology of most callitrichids, including all six *Callithrix* species (*C. aurita, C. flaviceps, C. geoffroyi, C. jacchus, C. penicillata*, and *C. kuhlii*) (Consortium, 2014; Harris et al., 2013).

**Fig. 1:**
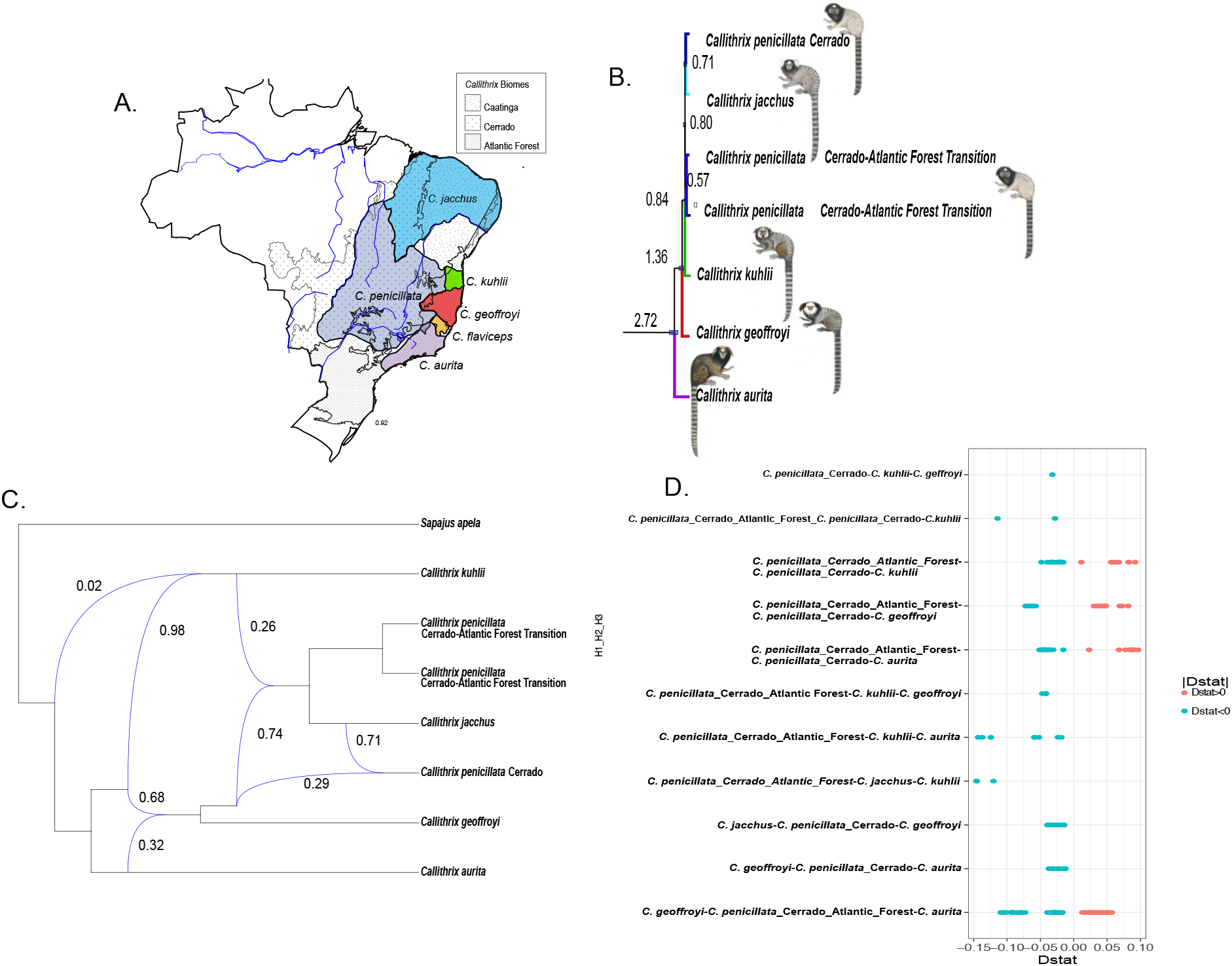
Phylogenomics, Divergence, and Ancient Introgression in *Callithrix*. (A) Natural geographic ranges of the six *Callithrix* species (color coded by species) and the three biomes in which these ranges occur. (B) Inset of *Callithrix* multispecies coalescent-based ASTRAL species tree and divergence time estimates as millions of years ago (MYA) shown at major nodes. Blue bars at nodes represent 95% divergence time confidence intervals. The full tree is shown in Supplementary Fig. S1A. (C) Phylonet reticulation network with blue lines representing interspecific gene flow between *Callithrix* species. Reticulations are shown as blue arrows with inheritance probability from the donor towards the recipient groups shown as percentages. (D) Significant D-statistics results for trio comparisons between individuals from different marmoset species. Along the y-axis are trios of “H1”, “H2”, and “H3”, whose order represents the assumed evolutionary relationship between the taxa as ((H1,H2),H3, outgroup). *Cebus imitator* represented the ancestral outgroup. *Callithrix* illustrations are by Stephan Nash and used with artist’s permission.

The *Callithrix* genus likely originated in the southeastern Brazilian Atlantic Forest and then migrated northward along the Brazilian coast and expanded into the semi-arid regions of northeastern and central Brazil (Buckner et al., 2015; Malukiewicz et al., 2021). *Callithrix aurita* along with *C. flaviceps* make up the *aurita* group of *Callithrix*, and the remaining species form the *jacchus* group. Based on mitogenomic data, the *aurita* group is basal in the *Callithrix* phylogeny (Malukiewicz et al., 2020) and weighs between 400g and 500g (Smith and Jungers, 1997; Malukiewicz et al., 2024). The *aurita* group species are found in the southern most part of the natural *Callithrix* distribution (Malukiewicz et al., 2020). It is known that *C. aurita* accesses tree exudates from sources which do not require them to gouge holes in tree bark (Martins and Setz, 2000; Malukiewicz et al., 2020), and this is also likely the case for *C. flaviceps*. The next species to diverge in the *Callithrix* phylogeny, *C. geoffroyi* and *C. kuhlii*, have natural distributions that occur north of that of *C. aurita*, weigh between approximately 350g and 400g (Smith and Jungers, 1997; Malukiewicz et al., 2020), and possess unique dental adaptations for extraction of tree exudates (Natori, 1986). Finally, *C. jacchus* and *C. penicillata*, the two most recent *Callithrix* species based on mitogenomic data (Malukiewicz et al., 2021), are found at the northern most portion of natural *Callithrix* distributions, weigh between approximately 250g to 350g, and are the most specialized marmosets for exudivory (Smith and Jungers, 1997; Malukiewicz et al., 2020, 2024). However, the specific genomic changes during *Callithrix* evolution that would have led to progressively smaller size and more extreme exudivory during evolution of the genus are not yet clear.

It is plausible that *Callithrix* migration from the Atlantic Forest and into Brazilian semi-arid biomes, the Caatinga and Cerrado, drove gradual reductions in body size and specialization for exudivory in this genus. The multi-layer tropical and humid Atlantic Forest naturally experiences little water stress (Brown and Prance, 1987; Filho and Filho, 2000). Habitats of the savanna-like Cerrrado vary from open grassland to denser woodland and experience less annual precipitation than the Atlantic Forest (Eiten, 1972). The lowest annual precipitation among Brazilian biomes occurs in the Caatinga, a seasonal dry tropical forest that receives most of its concentrated rainfall within three months but can also experience extended droughts (Sampaio, 1995; Chiang and Koutavas, 2004). It has also been shown that there is a negative relationship between average mammalian body mass and mean annual temperature (Hantak et al., 2021). *Callithrix* migration into more extreme habitats that show stark differences in vegetation and availability of nutritional resources relative to the Atlantic Forest may have driven changes in common callitrichid traits that led to the emergence of increasingly smaller and more exudate-focused marmoset species.

In this study, we employ low-coverage whole genome sequencing (lcWGS) from newly acquired genomic data from four *Callithrix* species (6 *C. aurita*, 5 *C. geoffroyi*, 7 *C. jacchus*, 8 *C. penicillata*), and obtained publicly available WGS data from a single *C. kuhlii* to refine our current understanding of the evolution of marmosets. We utilized the *Cebus imitator* genome (NCBI Accession GCA 001604975.1) as the reference assembly to minimize mapping bias between different marmoset species. The goal of this study was three-fold: (1) to reconstruct the evolutionary and demographic history of the *Callithrix* genus based on the first WGS genus-wide data available for *Callithrix* species; (2) to assess genetic diversity within and among *Callithrix* species; and (3) to identify the likely genetic basis of differences in body size and exudivory specializations within the *Callithrix* genus.

## Results and Discussion

### Phylogenomics the *Callithrix* Genus and Past Gene Flow

To construct a *Callithrix* species tree, we included at least one individual per species from those sampled in the wild with known provenance, with the exception of *C. jacchus* which was sampled in captivity. Due to the relatively wide natural geographical distribution and geographical structuring of *C. penicillata* by biome of origin from previous phylogenetic data (Malukiewicz et al., 2014, 2021), we included samples of this species from two different biogeographic regions (the Cerrado and the ecological transition between the Atlantic Forest and Cerrado). Quality-controlled and filtered Illumina FASTQ reads were aligned to the *Cebus imitator* genome assembly. Other WGS data were obtained from the NCBI database for *C. kuhlii* and 8 other publicly available primate genomes, which were also aligned to the *C. imitator* reference assembly (further described in Methods). All resulting BAM files from reference genome alignment were made into pseudohaplotype fasta files with ANGSD 0.930 (Korneliussen et al., 2014), which were then combined into a multiple species alignment (MSA). The MSA was divided into 250,000 base pair (bp) genomic windows that were filtered for ambiguous nucleotides (Ns), and maximum-likelihood (ML) trees were made from the filtered genomic windows with IQTREE v2.0.3 (Minh et al., 2020). We analyzed a total of 6824 genomic windows with ASTRAL 5.7.8 (Mirarab et al., 2014) to obtain a *Callithrix* multispecies coalescent tree (Fig. 1B and Supplementary Fig. S1A-B) with maximum support values (lpp=1.0) for all nodes except one (lpp=0.75 for the *C. jacchus*-*C. penicillata*-Cerrado node in Supplementary Fig. S1B). *Callithrix* divergence patterns within the topology shown in Fig. 1B are highly congruent with previously published phylogenies using mitogenonic (Malukiewicz et al., 2021), nuclear and mitochondrial (Reis et al., 2018), and phylogenomic data (Kuderna et al., 2023). Interestingly, *C. penicillata* is the only *Callithrix* species that naturally occurs in the Caatinga, the Cerrado, and the Atlantic Forest (Malukiewicz et al., 2020), and *C. penicillata* lineages show strong separation by geography and biome of origin in mitogenomic and nuclear phylogenies. The most basal *C. penicillata* clade originates from an area of ecological transition between the Cerrado and the Atlantic Forest, whereas younger clades originate from semi-arid regions in north Brazil (see Fig. 1A and Fig. 1B in this work and Fig. 2 in Malukiewicz et al. (2021)). *Callithrix jacchus* clusters with the youngest *C. penicillata* clades in the species tree in Fig. 1B and the mitogenomic phylogeny of (Malukiewicz et al., 2021). The species tree in Fig. 1B corroborates previous results that show a *Callithrix* origin in southeastern Brazil and then a northward migration by the genus (Buckner et al., 2015; Malukiewicz et al., 2021). Furthermore, based on phylogenetic results, we propose that *C. jacchus* originated in the semi-arid region of northeastern Brazil and then migrated to the Atlantic Forest of northeastern Brazil. However, this hypothesis will need to be tested in the future, ideally with expanded sampling of genetic material from *C. jacchus* from across all major geographic regions of the species’ natural distribution.

**Fig. 2:**
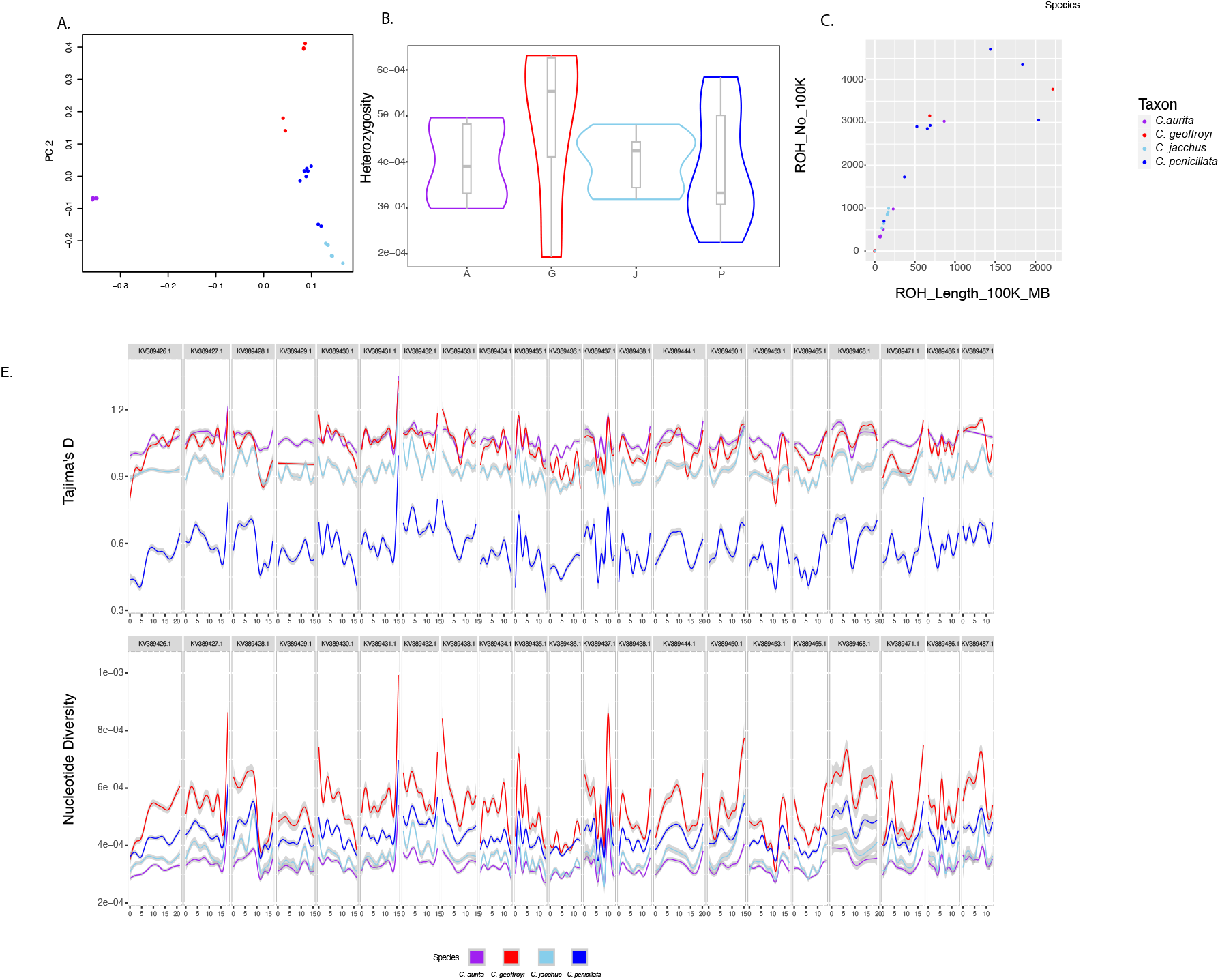
Population Genomics and Structure of *Callithrix* Species. (A) Principle components analysis of *Callithrix* species based on WGS, with each dot representing an individual marmoset. (B) Violin plots of average genome-wide heterozygosity for individual *Callithrix* species. The bottom box lines represent 25th percentiles, the mid-lines of boxes represent 50th percentiles/medians, and top box lines represent 75th percentiles. (C) Scatter plot of runs of homozygosity (ROH) for individuals in each *Callithrix* species. The x-axis shows the total length of ROHs for each individual in megabases (MB), and the y-axis shows the number of ROH tracts larger than 10 MB in each individual. (D) Species-specific smoothed Tajima’s D values (top) and nucleotide diversity (bottom) from genomic windows are shown along the 21 largest scaffolds in the *Cebus imitator* genome. Individual species follow the same color coding in each plot, and the figure legend seen on the right side show individual colors representing each species.

The divergences times we inferred for *Callithrix* species show that the genus diverged from the Cebidae clades of *Sapajus* and *Saimiri* approximately 13 million years ago (MYA), but *Callithrix* is a relatively recent radiation with *aurita* and *jacchus* groups diverging about 2.72 MYA, and divergence times between *jacchus* group species occurring between 0.71 and 1.36 MYA (Fig. 1B and Supplementary Fig. 1A). Previously published *Callithrix* divergence time estimates also show the *Callithrix* genus as a relatively young primate radiation. For example, Kuderna et al. (2023) estimates divergence times between *C. jacchus*-*C. kuhlii* -*C. geoffroyi* to have occurred between 0.46 and 0.66 MYA, while Reis et al. (2018) and Malukiewicz et al. (2021) placed the divergence between the *aurita* and *jacchus* groups at approximately 4 MYA and between *C. geoffroyi* and other *jacchus* group species divergences at less than 2.5 MYA. Deviation in estimates of divergence estimates between our and previously published works are likely explained by differences in methodology, time calibrations, and genomic/genetic markers.

The distribution of *Callithrix* ASTRAL quartet scores from the *Callithrix* multispecies coalescent does indicate a relatively high level of discord between the main species topology and the individual gene trees (Supplementary Fig. S1B). This discordance may be explained by the existence of a history of complex gene flow between *Callithrix* species, which we first investigated with Phylonet by reticulation network analysis based on 6975 genomic windows (Fig. 1C). We also looked for evidence of gene flow in the form of patterns of excessive derived allele sharing (D-statistic, Fig. 1D and Supplementary Table S1) between *Callithrix* lineages. Several results from these two approaches (Fig. 1C, Fig. 1D, and Supplementary Table S1) suggest the occurrence of gene flow between *Callithrix* species in line with the biogeographic and phylogenetic history of the genus as described above. For example, D-statistics indicated that *C. aurita* individuals share significantly more derived alleles with *C. geoffroyi* than with *C. penicillata*. The Phylonet approached indicated that the *jacchus* group retained 32% of the *C. aurita* genome due to gene flow between the most common recent ancestor (TMRCA) of the *jacchus* and *aurita* groups. Considering modern day geographical proximity between the natural ranges of *C. aurita* and *C. geoffroyi* as well as the phylogenetic proximity of these species, these results may point to past gene flow events between *aurita* and *jacchus* groups. As another example, the clade formed by *C. jacchus* and *C. penicillata* lineages may have experienced genetic introgression from *C. kuhlii* as well as TMRCA of *C. jacchus* and *C. penicillata*. Although the natural range of *C. kuhlii* lies in the northeast of Brazil in the Atlantic Forest, it abuts portions of the *C. penicillata* natural range that occur in the Caatinga (Fig. 1A). It is plausible that it was in this geographical region that TMRCA of *C. jacchus* and *C. penicillata* diverged from *C. kuhlii* and migrated out of the humid Atlantic Forest into more extreme semi-arid habitats while still maintaining gene flow with *C. kuhlii*.

In the *Callithrix* mitogenomic phylogeny (Malukiewicz et al., 2021), the *C. jacchus* clade is actually most phylogenetically proximate to *C. penicillata* lineages from the Caatinga. We would expect a similar phylogenomic result with the addition of *C. penicillata* lineages with known Caatinga provenance to our data, and well as evidence for past gene flow between *C. jacchus* and *C. penicillata* Caatinga lineages. We base this expectation on the result that *C. jacchus* also shows more significant sharing of derived alleles between *C. penicillata* Cerrado individuals than those of the *C. penicillata* Atlantic Forest-Cerrado transitional region (Fig. 1D), which is in line with the phylogenetic and geographic proximity of *C. jacchus* with *C. penicillata* from the Cerrado. Additionally, the Cerrado *C. penicillata* lineage retained 71% of the *C. jacchus* genome while the remainder came from the ancestor of *C. jacchus*/*C. penicillata* (Fig. 1C). With the inclusion of *C. penicillata* individuals with known provenance from the Caatinga, we would expect Phylonet and D-statistic results to point to the occurrence of gene flow between *C. jacchus* and *C. penicillata* Caatinga lineages.

Significant D-statistic results indicate the occurrence of secondary gene flow (transfer of genes between previously geographically separated populations that have already started to diverge) between *C. penicillata* and other *Callithrix* species (Fig. 1D, Supplementary Table S1). *Callithrix penicillata* has the largest natural distribution of all *Callithrix* species, and is the only *Callithrix* species whose natural distribution abuts the natural distributions of all other *Callithrix* species. D-statistic results indicate significant sharing of derived alleles between *C. aurita* and *C. geoffroyi*, respectively, with *C. penicillata*. After *C. penicillata* initially diverged within the semi-arid portions of northeastern Brazil, it likely spread out into other portions of Brazil. The spread of *C. penicillata* likely included a downward migration into southeastern Brazil where *C. penicillata* eventually encountered *C. aurita* and *C. geoffroyi* at the limits of their respective natural distributions. Upon these encounters, secondary gene flow may have occurred between each respective species pair, and natural hybrids between *C. penicillata* and *C. aurita* have been observed at natural geographic borders of these two species (Passamani, 2018).

### *Callithrix* Population Genomics

In the Principle Components Analysis (PCA) plot in Fig. 2A, *C. aurita* is clearly discriminated into a tight cluster along PC1 away from the *jacchus* group species. Then, we see discrimination of the *jacchus* group species along PC2 in the PCA plot into species-specific cluster. Within the *C. geoffroyi* cluster, samples which grouped in the upper right corner were collected in the eastern-most part of the species’ range in the state of Espírito Santo and those grouping closer to the *C. penicillata* cluster were collected in northeastern Minas Gerais state, in the western-most part of the *C. geoffroyi* range. These geographic extremes of the *C. geoffroyi* range occur in two different biomes, with the western-most portion occurring within the Cerrado and the eastern-most portion occurring within the Atlantic Forest. Within the *C. penicillata* cluster, we see clear separation of two individuals away from the main cluster and more closely with the *C. jacchus* cluster. The original provenance of these two individuals, which were sampled in captivity, is unknown.

To measure *Callithrix* genetic diversity, we calculated average per site heterozygostiy (Fig. 2B), runs of homozygosity (ROH) (Fig. 2C), genome-wide Tajima’s D (Fig. 2D top and Supplementary Fig. S2), and genome-wide nucleotide diversity (Fig. 2D bottom and Supplementary Fig. S3) in *C. aurita, C. geoffroyi, C. jacchus*, and *C. penicillata. Callithrix kuhlii* was excluded due to lack of population level genomic data. In order to reduce bias in sequencing coverage, WGS data from four high coverage samples was down-sampled to 1.5x to reflect average coverage of our sample set (see Methods). *Callithrix geoffroyi* showed the highest median genome-wide heterozygosity, while *Callithrix penicillata* showed the lowest median genome-wide heterozygosity values (Fig. 2B). *Callithrix aurita* is considered one of the most endangered primates, with about 10,000-11,000 adult animals remaining in the wild due to a population decline of >50% between 2000-2018 attributed to heavy habitat loss (>43%) from land development, logging and wood harvesting, invasive species and hybridization (Malukiewicz et al., 2021; Melo et al., 2015). However, the *C. aurita* heterozygosity range was still within those of the other three *Callithrix* species (Fig. 2C). Most individual *Callithrix* total ROH lengths were less than 500 MB, but *C. penicillata* individuals tended to have the highest ROH length out of all *Callithrix* species (Fig. 2C). ROHs also suggest minimal recent inbreeding in *C. aurita*, as the species overall manifested relatively low total ROH length relative to the other species (Fig. 2C).

*Callithrix penicillata* showed the lowest Tajima’s D values relative to the other *Callithrix* species, and the lower part of the interquantile range fell into negative values (Fig. 2D and Supplementary Fig. S2), which can indicate population expansion or recent selective sweeps. As *C. penicillata* possesses the widest *Callithrix* natural geographic distribution, its genome-wide pattern of Tajima’s D values may reflect recent population expansion throughout northeastern, central, and southeastern Brazil. *Callithrix aurita* showed relatively the highest Tajima’s D values across most of the 21 longest contigs of the *C. imitator* reference assembly (Fig. 2D), and the highest median genome-wide Tajima’s D value (Supplementary Fig. S2). Positive Tajima’s D values could indicate balancing selection or sudden population contraction. The latter may apply for *C. aurita*, due to the species’ highly endangered status (Malukiewicz et al., 2021). This species also showed the lowest nucleotide diversity values in the 21 longest *C. imitator* contigs (Fig. 2D), but its median genome-wide nucleotide diversity values were similar to *C. penicillata* and *C. jacchus* (Supplementary Figure S3). When considering genetic diversity as a proxy for the adaptive potential of a population (Lande and Shannon, 1996), it is plausible that the *C. aurita* population potential, despite the species’ endangered status (Malukiewicz et al., 2021), is within the potential of other *Callithrix* species. Relative to *C. geoffroyi*, lower levels of genetic diversity for *C. jacchus* and *C. penicillata* (Fig. 2D and Supplementary Fig. S3) maybe due to less accumulated genetic diversity in the former species given their more recent divergence times among *Callithrix* species.

We also conducted genome-wide Fst scans in 50000 base pair windows in 10000 base pair steps. We see that the highest genome-wide Fst average is found between the two most phylogenetically diverged species pairs, that being *C. aurita*-*C. jacchus* and *C. aurita*-*C. penicillata* (Supplementary Table S2). Then the lowest genome-wide average was found for the most recently phylogenetically diverged pair, *C. jacchus* and *C. penicillata* (Supplementary Table S2). A similar pattern is observed when we look at pairwise Fst values across genomic windows mapped to the 21 largest scaffolds in the *Cebus imitator* reference genome (Supplementary Fig. S4).

### Functional Enrichment Analysis of *Callithrix* Candidate

#### Positive Selection Genes

To identify potential candidate genes targeted by selection during *Callithrix* speciation, we conducted genome-wide scans of the population branch statistic (PBS) (Fig. 3, Supplementary Tables S3-S14), which were included in the same Fst analysis as described above. The PBS approach can identify targets of recent selection in a focal population relative to differences in allele frequencies compared against two outgroups. The PBS approach has been found to outperform Fst at identifying local sweeps, and shows greater specificity in detecting population-specific positive selection (Shpak et al., 2024). We considered outlier genomic windows as those possessing the top 99.9th percentile of genome-wide PBS values, and genes present within each outlier window were considered to represent candidate genes under positive selection. Then, we focused functional enrichment analysis on PBS candidate selection genes, which was conducted in g:GOSt with the reference organism set to *C. jacchus*. The total number of candidate genes among various PBS trios included in the g:GOSt analyses ranged from 49 to 104 genes (Supplementary Tables S3-S14).

**Fig. 3:**
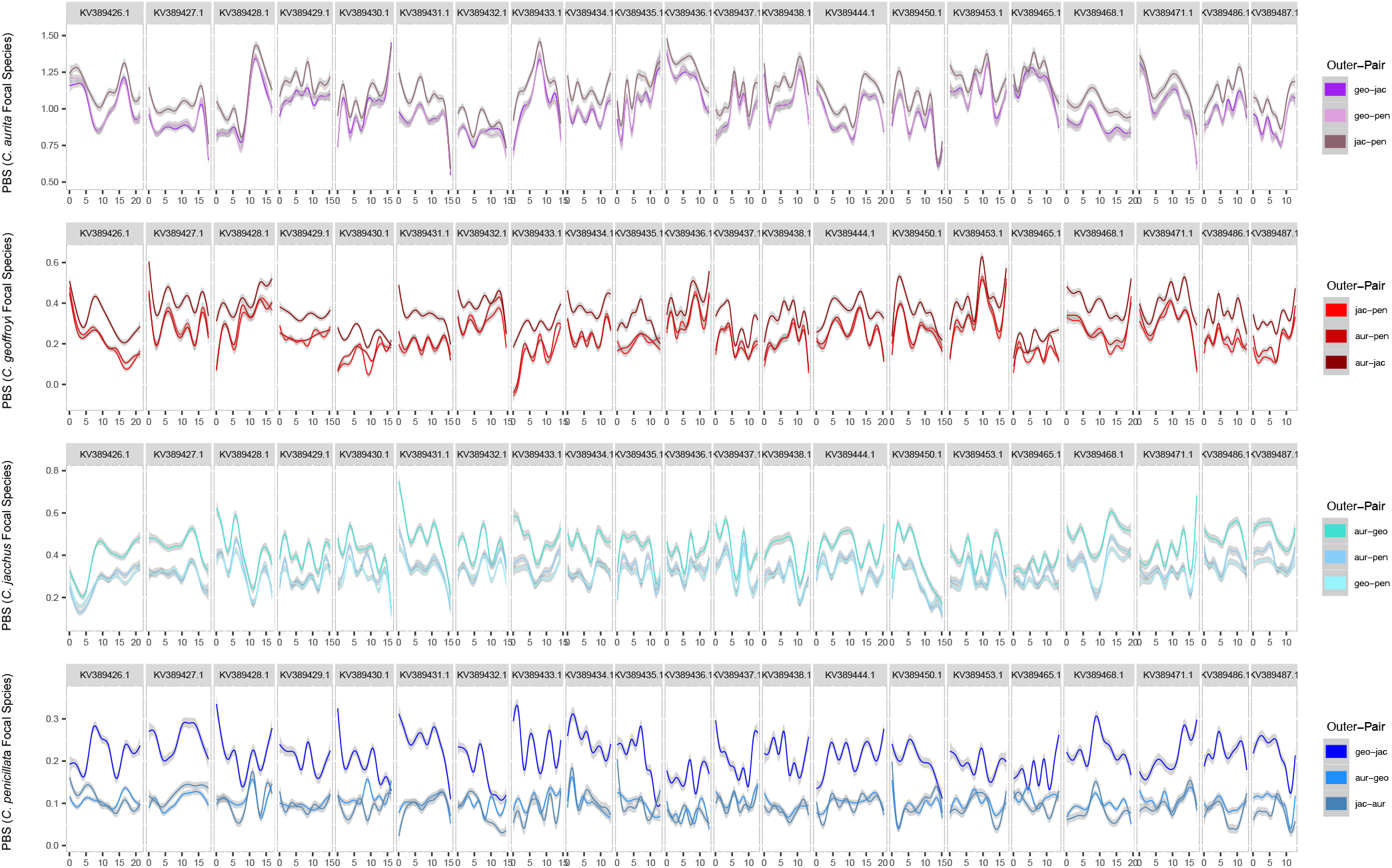
Population branch statistic (PBS) values for genome-wide genomic windows between all possible trios of *Callithrix* species. Each plot panel features comparisons between a unique “focal” species and pairs of two “outgroup” species. Smoothed PBS values from genomic windows are shown along the 21 largest scaffolds in the *Cebus imitator* genome. Abbreviations in each legend correspond to species names as follows: aur-*C. aurita*, geo-*C. geoffroyi*, jac-*C. jacchus*, pen-*C. penicillata*.

Next, we used the Cytoscape EnrichmentMap app to find possible overlaps of significant functionally enriched GO terms between PBS trios with the same focal species. In all PBS trios where *C. aurita* and *C. jacchus* were the respective focal species (GO enrichment results for individual trios Supplementary Tables S15-S20), we found overlapping significant functional GO terms related to anatomical development, stress and stimulus response, and regulation of biological processes (Supplementary Fig. S5 and S6). We found no overlapping significant GO terms in PBS trios where *C. geoffroyi* was the focal species (GO enrichment results for individual trios Supplementary Tables S21-S23), but significant GO terms for individual trios were related to collagen fibrils, and estrogen receptor activity. We found no overlapping significant GO terms in PBS trios where *C. penicillata* was the focal species (Supplementary Tables S24-S26, Supplementary Fig. S7), but for individual trios significant GO terms included cell morphogenesis, cellular junctions and synapses, and regulation related to cytokines.

#### Selection in Candidate Speciation Genes Related to Morphological Traits

From the PBS analyses described above, we identified several candidate genes under selection for craniofacial development and morphology in the main *Callithrix* species included in our study (Fig 4). Morphological differences in the mandibular and zygomatic cranial portions between *C. aurita* and *C. jacchus*/*C. penicillata* have been attributed to differences in dietary specialization between these taxa (de Souza, 2016; Malukiewicz et al., 2024). *Callithrix jacchus* and *C. penicillata*, are also the most morphologically specialized for an exudivorous diet and to gouge holes in trees to access tree gums (Rylands and Faria, 1993). On the other hand, *C. aurita* only accesses exudates opportunistically without gouging and is relatively less morphologically specialized for exudivory (Rylands and Faria, 1993). For *C. aurita*, morphology related candidate genes include *SZT2* (Naseer et al., 2018), *HDAC9* (Hirsch et al., 2022), and *ARHGAP29* (Letra et al., 2014). Candidate genes for *C. jacchus* included *IGFBP5* (regulator of craniofacial skeletogenesis (Amaar et al., 2006)) and *TCOF1*, which was also identified as a candidate for positive selection in *C. penicillata*. Thus, *TCOF1* may be of particular importance in the evolution of craniofacial morphology and gumnivory in *Callithrix*. In humans, *TCOF1* mutations cause Treacher Collins syndrome, which is characterized by downward-slanting palpebral fissures, mandibular hypoplasia, and zygomatic hypoplasia (Dixon et al., 2006; McElrath and Winters, 2023). The *jacchus* group of marmosets shows greater protrusion of the upper and lower jaws and more compressed brain-case relative to the *aurita* group of *Callithrix* marmosets (Souza, 2016). We suggest that *TCOF1* should be further studied as a candidate gene associated with craniofacial morphological differences between marmoset species.

**Fig. 4:**
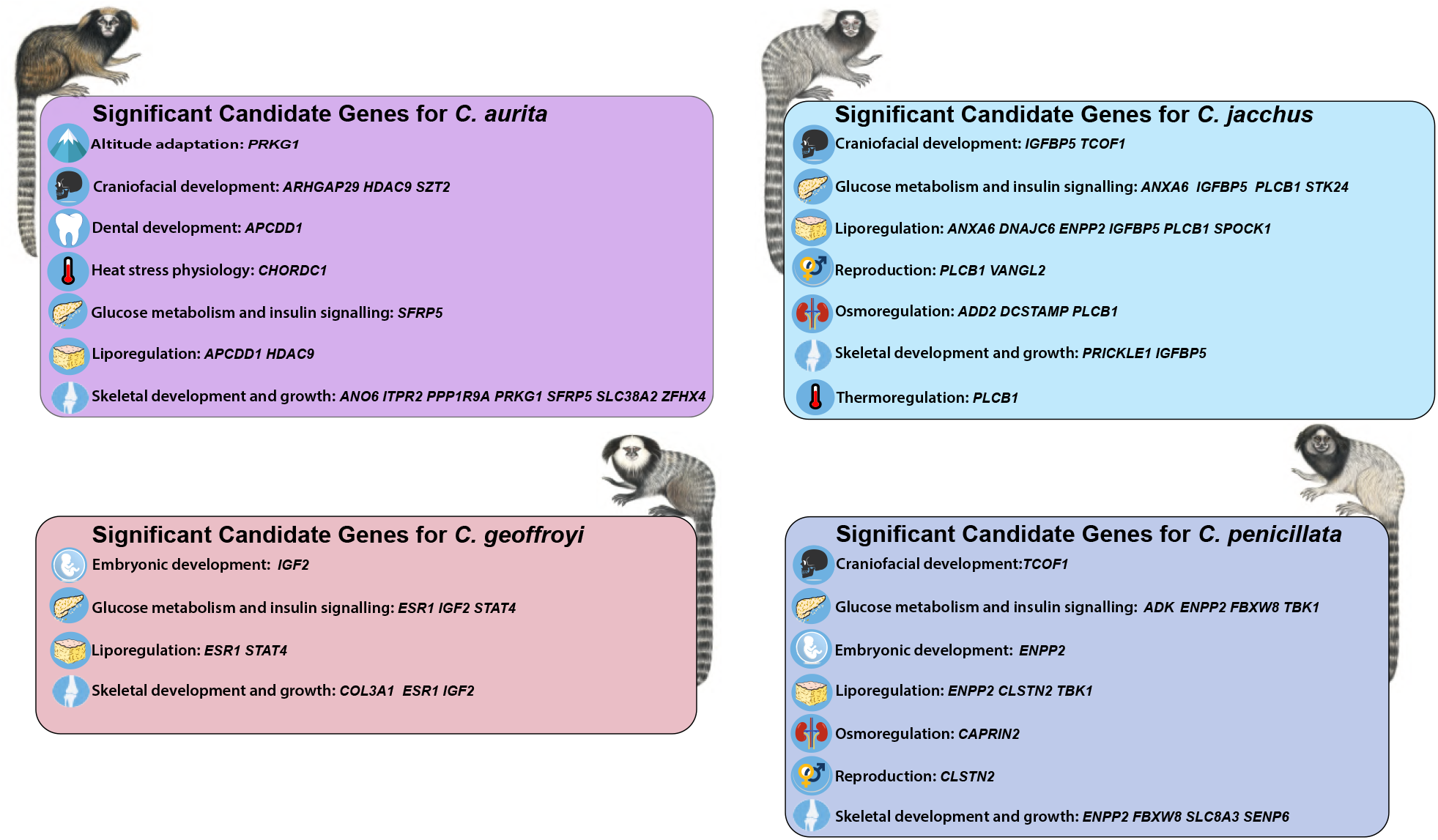
Examples of *Callithrix* candidate genes putatively under positive selection. For each studied *Callithrix* species, examples are shown of candidate genes whose biological function has been previously shown to be associated with metabolism, growth/development, and homeostasis. *Callithrix* illustrations by Stephan Nash and used with artist’s permission.

Miniaturization of body size in marmosets and other callitrichids is achieved, in part, by the slowing down of pre-natal growth rates (Marroig and Cheverud, 2009; Soman et al., 2024), which is unique among mammals (Marroig and Cheverud, 2009). From the PBS analysis, we identified several candidate genes that may have influenced body size in *Callithrix* (Fig. 4). In *C. aurita*, these genes included *ANO6* (gene knockout causes reduced skeleton size (Ehlen et al., 2013)), *SLC38A2* (controls fetal development and postnatalbone formation (Matoba et al., 2019; Mandò et al., 2013; Shen et al., 2022)), *ZFHX4* (coordinates endochondral ossification and longitudinal bone growth (Nakamura et al., 2021; Fontana et al., 2021)), *ITPR2* (causes decreased limb length and short stature (Moreno-Reyes et al., 1998; He et al., 2019)), and *PPP1R9A* (associated with embryonic growth and birth weight (Zaitoun and Khatib, 2008; Tunster et al., 2013; Zhang et al., 2014)). *SFRP5*, as another candidate gene in *C. aurita*, may function as a potential negative regulator of bone mass (Chen et al., 2016). In *C. jacchus*, candidate genes included *PRICKLE1* (stunts limb growth (Yang et al., 2013); contributes to bone development (Wan and Szabo-Rogers, 2020)), *ENPP2* (involved in osteoblast differentiation and mineralization (Tourkova et al., 2024)), *IGFBP5* (stimulates bone formation (Amaar et al., 2006)), and *DCSTAMP* (plays critical roles in osteoclast fusion and master regulator of osteoclastogenesis, (Zhang et al., 2014; Chiu and Ritchlin, 2016; Yagi et al., 2006)). For *C. penicillata*, we identified the aforementioned *ENPP2* and *FBXW8* (gene knockout stunts growth (Tsutsumi et al., 2008)), as well as *SLC8A3* (involved in mineralization of osteoblasts (Sheikh et al., 2024)) and *SENP67* (maintains osteochondroprogenitor homeostasis (Li et al., 2018)) as under putative positive selection. In *C. geoffroyi, IGF2* (influences growth and adipocity (Kadakia and Josefson, 2016)) and *COL3A1* (role in development of trabecular bone (Volk et al., 2014)) may be under positive selection as well. The *C. penicillata* candidate genes *ENPP2* (involved in early embryogenesis signaling (Frisca et al., 2016)) and *IGF2* (influences prenatal and postnatal growth (Roberts et al., 2008; Smith et al., 2006; Begemann et al., 2015)) may have pleiotropic effects also on embryonic development. The gene *ESR1*, which is involved with trabecular bone formation (Windahl et al., 2013), and affects female bone density (Mondockova et al., 2018; Krela-Kaźmierczak et al., 2019)), was found to be a candidate gene in *C. geoffroyi*. In *C. jacchus*, behaviorally subordinate females experience long periods without ovulation and low circulating levels of reproductive hormones, including estrogen (Saltzman et al., 2018). However, female *C. jacchus* do not show estrogen-depletion bone loss as females of almost all other mammalian species do (Saltzman et al., 2018). Estrogen receptors, as suggested by our results for *ESR1* in *C. geoffroyi*, may be part of unique adaptations marmosets have developed to avoid estrogen-depletion bone loss.

#### Positive Selection Related to Stress Response, Homeostasis, Glucose Metabolism, and Fat Metabolism

From the PBS analysis above, we identified several candidate genes under putative positive selection whose biological function has been previously shown to be related to stress response, homeostasis, glucose metabolism, and fat metabolism (Fig 4). We were interested in how these genes may be related to challenges that *Callithrix* marmosets face in the different biomes that are part of the natural *Callithrix* distribution. For example, *C. aurita* occupies the mountainous regions of the southeastern Atlantic Forest (80-1300 meters (Melo et al., 2021)) where average temperatures vary from 17°C at high altitudes to 22°C degrees at low altitudes, as in the Serra do Mar mountains (Joly et al., 2012). We identified two candidate genes in *C. aurita* implicated in thermal tolerance, *PRKG1* and *CHORDC1* (Cardona et al., 2014; Dou et al., 2020, Dou et al., 2021), which may help this species tolerate variable temperatures at different altitudes. On the other hand, *C. jacchus* faces a wider range of average annual temperatures in the Caatinga (e.g. in the state of Ceará the average temperatures vary from 22.4°C ± 0.38 to 33.5°C± 0.69 (Campos and De Andrade, 2021). The *C. jacchus* candidate selection gene *PLCB1* for heat stress regulation (Aboul-Naga et al., 2022) may help thermoregulation in this species at different times of the year.

For mammals native to arid environments, it is expected that they would experience selection for adaptations related to fat metabolism, insulin signaling and water retention (Rocha et al., 2021), and we extend this expectation to Brazilian semi-arid environments as well. For osmoregulation, we identified *CAPRIN2* as a candidate gene in *C. penicillata* (involved with physiological osmotic stress response, (Konopacka et al., 2015)) and in *C. jacchus*, we identified *ADD2* (associated with blood pressure and hypertension via renal tubule sodium reabsorption (Zhang et al., 2019)). For *C. jacchus*, candidate genes whose functional role involves liporegulation include (1) *ANXA6* (associated with cholesterol and modulation of triglyceride accumulation and storage (Grewal et al., 2010; Cairns et al., 2018; Krautbauer et al., 2017)); hepatic glucose metabolism (Cairns et al., 2018); (2) *PLCB1* (adipocyte proliferation (Liang et al., 2024)); (3) *IGFBP5* (cellular lipid accumulation of lipids in muscle cells (Xiang et al., 2019; Xiao et al., 2020)); (4) *SPOCK1* (differentiates adipose tissue (Alshargabi et al., 2020)); (5) *ENPP2* (involved with fatty liver disease and liver lipid remodeling (Brandon et al., 2019)); and (6) *DNAJC6* (involved with adipogenesis with energy metabolism (Son and Lee, 2023); early-onset obesity (Vauthier et al., 2012)). For *C. penicillata*, candidate genes involved in liporegulation include *ENPP2, CLSTN2* (involved with HDŁmediated cholesterol efflux capacity (Schachtl-Riess et al., 2023)), and *TBK1* (involved in energy homeostasis and in adipose tissue (Zhao et al., 2018; Zhang et al., 2020)). Interestingly, we found candidate genes related to temperature-sensitive liporegulation in *C. aurita*, including *HDAC9* (involved with temperature-sensitive adipogenic differentiation (Chatterjee et al., 2013; Goo et al., 2022; Ahmadieh et al., 2023) and *APCDD1* (adipocyte differentiation (Yiew et al., 2017)), which may be important for altitude related thermal tolerance.

Candidate genes involved in insulin signaling and glucose metabolism in *C. jacchus* include *ANXA6* (hepatic glucose metabolism (Cairns et al., 2018)), *PLCB1* (pancreatic beta-cell insulin secretion (Hwang et al., 2019)), *STK24* (inhibition of insulin signaling (Iglesias et al., 2017)), and *IGFBP5* (insulin sensitivity (Xiao et al., 2020); pancreatic beta-cell growth (Gleason et al., 2010)). For *C. penicillata*, candidate genes involved with insulin signaling included *FBXW8* (restriction of insulin signaling pathways (Mieulet and Lamb, 2008; Kim et al., 2012)), *TBK1* (regeneration of pancreatic beta-cells, (Jia et al., 2020)) and *ADK* (regulation of diet-related insulin resistance (Xu et al., 2019); inhibition of beta-cell proliferation (Chen et al., 2024)). We also identified candidate genes in *C. geoffroyi* that included *ESR1* (gluconeogenesis (Qiu et al., 2017); liver carbohydrate metabolism (Khristi et al., 2019)), and the aforementioned *IGF2* (regulation of pancreatic growth and function (Hammerle et al., 2020)), and *STAT4* (glucose intolerance and insulin resistance Dobrian et al. (2017)). *SFRP5*, which was mentioned above in *C. aurita*, may be a pleiotropic candidate selection gene in this species that is also involved in insulin sensitization and anti-inflammation (Ouchi et al., 2010; Rebuffat et al., 2013; Hu et al., 2013; Guan et al., 2016).

All *Callithrix* species eat tree gums to some extent and these various candidate selection genes may be important to glucose metabolism related to dietary exploitation of oligosaccharides. Particularly, for the species which are highly specialized for gumnivory, tree gum is a key seasonal nutritional resource during the driest part of the year in Brazilian semi-arid regions when food supply is limited. Insulin resistance is a plausible mechanism that marmosets have evolved to account for food scarcity, and natural insulin resistance without diabetic pathology is a phenotype seen in mammals that may be beneficial under food scarcity to evade starvation and reduce energetic demands (Rocha et al., 2021; Houser et al., 2013; Rigano et al., 2016; Blanco et al., 2023). Perhaps some the putative positive selection candidate genes we have identified that are involved in insulin signaling and glucose metabolism have allowed marmosets to evolve some form of insulin resistance. However, we put this idea forth as a speculative hypothesis that would need further scientific investigation.

## Conclusion

Currently, the availability and development of genomic resources for marmosets and callitrichids in general are focused mainly on *C. jacchus* due to its importance as an emerging biomedical model. This study greatly expands population level genomic data available for four out of the six *Callithrix* species and sheds light on the genomic basis of the evolutionary changes likely linked to derived small body size and exudivory in marmosets. Our results indicate that *C. jacchus* and *C. penicillata* may have experienced changes in genes linked to fat and glucose metabolism which were not found in the most recent common *Callithrix* ancestor. These finding suggest that husbandry practices of *C. aurita* and *C. jacchus*/*C. penicillata* differ as the former is an opportunistic exudivore while the latter are highly specialized exudivores. As a follow up to the genomic scans presented here, future genomic studies of marmosets should combine higher-coverage genomic, transcriptomic, phenotypic, and climatic data in a larger sample of individuals per species across the entire *Callithrix* native range. This approach will allow for linking data on genetic variation in marmoset species with functional and phenotypic data across the different environmental niches occupied by marmosets. As three out of the six *Callithrix* species are endangered, knowing the range of *Callithrix* genetic variation, particularly adaptive genetic variation, will help guide decisions about how to preserve the adaptive potential of marmoset species. Further, as captivity of marmosets is important for biomedical and conservation reasons, our data contribute towards biological understanding of differences among marmoset species. We highly urge captive facilities to incorporate this new knowledge towards improving husbandry of captive marmosets to increase their welfare under captive conditions.

## Methods

### Sample Populations

Sampling of marmosets occurred in Brazil between 2011 and 2016. In 2011, skin tissue was collected from two *C. penicillata* individuals that were captured in Brasília, Federal District. Between 2010 and 2016, skin tissue was collected from: (1) wild marmosets in Minas Gerais and and Espírito Santo states, as well as the Brazilian Federal District; (2) captive-born, wild-caught, and confiscated marmosets housed at the Guarulhos Municipal Zoo, Guarulhos, São Paulo state, CEMAFAUNA (Centro de Manejo de Fauna da Caatinga), Petrolina, Pernambuco state, and Centro de Primatologia do Rio de Janeiro (CPRJ), Guapimirim, Rio de Janeiro state; (3) a wild group of *C. aurita* from Natividade, Rio de Janeiro state that was caught and housed at CPRJ. Sampling consisted of a total of newly sequenced 26 *Callithrix* individuals (Supplementary Table S27), and one previously sequenced *C. kuhlii* individual (NCBI Bioproject SRX834392). Marmoset capture and sampling methodology has been described elsewhere (Malukiewicz et al., 2014). All individuals were allowed to recover after sample collection, and wild marmosets were released at their point of capture. Specimens were classified phenotypically as pure *C. aurita, C. geoffroyi, C. jacchus* and *C. penicillata* based on previously available descriptions (Hershkovitz, 1977; Fuzessy et al., 2014; Malukiewicz et al., 2014; Carvalho, 2015; Malukiewicz et al., 2024).

### Laboratory Procedures and Sequencing

DNA from skin samples was extracted using a standard proteinase K/phenol/chloroform protocol (Sambrook and Russell, 2001). Buffers used for extraction, precipitation and elution of DNA are listed elsewhere (Malukiewicz et al., 2014). DNA from marmosets collected in Brasília, Federal District were extracted at Northern State Fluminense University, Rio de Janeiro State, and then exported to ASU (CITES permit #11BR007015/DF). DNA from all other individuals was extracted at the Federal University of Viçosa (UFV), Viçosa, Minas Gerais, Brazil. Genetic samples collected since 2015 have been registered in the Brazilian CGen SISGEN database (Supplementary Table S27).

Low-to-medium coverage (1x-15x) whole genome sequencing was carried out for each sampled individual. WGS sequencing libraries were prepared at UFV and ASU with Illumina Nextera DNA Flex Library Prep Kits (catalog #20018704) following manufacturer’s instructions. Individual libraries were barcoded with Illumina Nextera DNA CD Indexes (catalog # 20018707), and pooled in equimolar amounts and sequenced on an Illumina NextSeq sequencer using v2 chemistry for 2 x 150 cycles at the ASU Genomic Core Facilities, Tempe, AZ, USA. Newly sequenced WGS data have been deposited in ENA (Bioproject Accession Number PRJEB88615) and NCBI (Bioproject Accession Number PRJNA1258696).

### Reference Genome Mapping, Variant Detection, and Filtering

Raw Illumina sequencing reads were initially checked for quality with FastQC v0.11.7 (https://www.bioinformatics.babraham.ac.uk/projects/fastqc/). Then leading low quality or N bases were trimmed from reads with Trimmomatic v0.36 (Bolger et al., 2014) and Illumina adapter sequences removed from FASTQ reads with Trim Galore! v0.4.3 (http://www.bioinformatics.babraham.ac.uk/projects/trim_galore/). For reference genome alignment, we followed a strategy to reduce reference mapping bias (van der Valk et al., 2019), and we aligned filtered FASTQ reads to the *Cebus imitator* genome assembly (NCBI Accession GCA 001604975.1), one of the closest available outgroups for marmosets. All genomic alignments from each sampled individual were made with the BWA-MEM v0.7.10 algorithm (Li and Durbin, 2009) with default settings. PICARD 2.18.27 (http://broadinstitute.github.io/picard) sorted resulting BAM files, and SAMTOOLS 1.10 (Danecek et al., 2021) filtered reads from BAM files with flag 3844. This flag removes reads that were unmapped, not in the primary alignment, were PCR or optical duplicates, or a supplementary alignment. Duplicate reads in individual files were marked and removed with the PICARD MarkDuplicates tools. Finally, local realignment around indels in individual BAM files was conducted with the GATK 3.7 IndelRealigner tool (van der Auwera and O’Conner, 2020). After the realignment step, averaged genomic-wide coverage for sequencing reads was calculated as the average of the ‘mean-depth’ column from the output produced with the SAMTOOLS coverage command. The number of mapped reads for each set of realigned sequencing reads was calculated from output the SAMTOOLS flagstat command.

### *Callithrix* Phylogeny and Divergence Dating

To construct the *Callithrix* species tree, we included at least one individual per species. Due to the relatively wide natural geographical distribution of *C. penicillata* and geographical structuring of this species by biome of origin from previous phylogenetic data (Malukiewicz et al., 2014, 2021), we included samples of this species from two different biogeographic regions (the Cerrado and the ecological transition between the Atlantic Forest and Cerrado). We prioritized individuals with the highest average coverage that represented at least one individual per *Callithrix* species (BJT6-*C. aurita*; BJT169-*C. geoffroyi*; cpe025, BJT8 and BJT40-*C. penicillata*; BJT157-*C. jacchus*). Whole genome sequencing data were obtained for *C. kuhlii* from NCBI under accession number SRX834392. As phylogenetic outgroups, we utilized the following publicly available primate genomes: (1) *Carlito syrichta* (Tarsius-syrichta-2.0.1.dna-rm.toplevel.fa.gz from the Ensembl database); (2) *Sapajus apella* (NCBI Accession GCF-009761245.1-GSC-monkey-1.0-genomic.fna); (3) *Gorilla gorilla* (Gorilla-gorilla.gorGor4.dna-rm.toplevel.fa.gz from the Ensembl database); (4) *Pan troglodytes* (Pan-troglodytes.Pan-tro-3.0.dna-rm.toplevel.fa.gz from the Ensembl database); (5) *Papio anubis* (Papio-anubis.Panubis1.0.dna-rm.toplevel.fa.gz downloaded from the Ensembl database); (6) *Saimiri boliviensis boliviensis* (Saimiri-boliviensis-boliviensis.SaiBol1.0.dna-rm.toplevel.fa.gz (downloaded from the Ensembl database); (7) *Plecturocebus donacophilus* (NCBI Accession GCA-004027715.1); and (8) *Pithecia pithecia* (NCBI accession number GCA-023779675.1). Genomic data from the above outgroups were aligned to *Cebus imitator* reference genome with MINIMAP2 2.17 (Li et al., 2018) and the -a flag to obtain BAM files.

Then using ANGSD 0.930 (Korneliussen et al., 2014), we made pseudohaplotype fasta files for each sample using BAM files resulting from the BWA-MEM and MINIMAP2 alignment. ANGSD was run with the input BAM files, the reference genome set to that of *Cebus imitator*, and parameters set to: -dofasta 1 - doCounts 1 -P 4. Using custom scripts, multiple species alignments (MSAs) of *Callithrix* and outgroup samples were created for each scaffold for the *C. imitator* genomic assembly. Only a subset of MSAs from the 500 largest *Cebus imitator* scaffolds by length were included in construction of the *Callithrix* species tree, as outgroups and marmoset samples tended to not map to smaller size *Cebus imitator* scaffolds. Resulting FASTA files were repeat masked with REPEATMASKER v4.1.4 (http://www.repeatmasker.org) using standard program settings. Using custom scripts, each MSA was divided into 250000 base pair genomic windows which were then filtered for N within the sequences.

Maximum Likelihood (ML) trees were made from filtered genomic windows with IQTREE v2.0.3 (Minh et al., 2020) using the command “iqtree -s example.phy -B 1000 -alrt 1000.” This command combines ModelFinder (Kalyaanamoorthy et al., 2017) to automatically find the best evolutionary model for each tree, conduct the ML tree search, and carry out the SH-aLRT test and ultrafast bootstrapping with 1000 replicates per tree search. All resulting ML trees were combined into a single file as input for ASTRAL 5.7.8 (Mirarab et al., 2014) to make a final species tree for *Callithrix*. ASTRAL was run with default settings.

Divergence dating of the *Callithrix* species tree resulting from the ASTRAL analysis was performed with the Reltime-ML option in MEGA11 (Tamura et al., 2021) following the protocol of Mello (2018). Dating calibrations were set to a divergence time of 41 million years ago (MYA) with a minimum only bound between Anthropoidea and Tarsiiformes, divergence time with uniform distribution of 33.4-56.035 MYA between the Catarrhini and Platyrrhini, divergence time with uniform distribution of 25.193-33.4 MYA between Cercopithecoidea and Hominoidea, a uniform distribution of divergence time of 7.5-15 MYA between Gorillini and Homini, a minimum bound divergence time of 13.363 MYA between Pitheciidae-(Aotidae+Atelidae+Callitrichidae+Cebidae), a uniform distribution divergence time of 13.183-34.5 MYA between Callitrichidae-Cebidae, and a uniform distribution divergence time of 13.032-34.5 MYA between Cebinae-Saimirinae (de Vries and Beck, 2023). *Carlito syrichta* was set as the outgroup in the Reltime-ML divergence analysis. Reltime-ML options were set to default which included the general time reversible model, uniform rates among sites, and local clock type.

### Past *Callithrix* Hybridization

To find evidence of past introgression between *Callithrix* species, we used PHYLONET 3.8.2 (Wen et al., 2018) to reconstruct a *Callithrix* reticulate network using *Sapajus apella* as an outgroup. Each *Callithrix* species was represented by a single individual which we subsampled from our full panel of samples, prioritizing samples with known provenance. We inferred the *Callithrix* network with the InferNetwork MPL command, which uses maximum pseudo-likelihood (MPL) and gene trees as input. Therefore, ML gene trees were generated with IQTREE as previously described the ASTRAL *Callithrix* species tree. IQTREE trees were then rooted with *Sapajus apella* using a custom python script based on the tree.reroot at edge command of the Dendropy Python library (Sukumaran and Holder, 2010). We then ran the InferNetwork MPL command with default settings by configuring a NEXUS input file. Individual ML trees were aggregated together into a NEXUS format file, which contained a Phylonet block in the format of “BEGIN PHYLONET; InferNetwork MPL (all) X;END;.” X within the block represented the number of reticulate events to reconstruct within the resulting network, which we varied from 0-4. We compared the top inferred networks between all runs representing a different number of possible reticulations, and the optimal number of reticulation events was estimated by searching the global optimum of the pseudolikelihood global optimums (Cao et al., 2019). Then we took the inferred network with the least negative MPL score among these top five candidate networks to represent the most likely *Callithrix* reticulation network.

As a complimentary test of ancient admixture, we used the ANGSD doAbbababa command line to calculate the D-statistic, or the so-called ABBABABA test, with the following set of flags “-uniqueOnly 1 -remove bads 1 -only proper pairs 1 -trim 0 -C 50 - baq 1 -P 10 -minMapQ 25 -minQ 25 -setMinDepth 1 -setMaxDepth 50.” We carried out the ABBABABA test for all possible trios of H1, H2, H3, and H4 (with *Cebus imitator* as the representative of the ancestral H4 variant) in which H1 and H2 did not represent sister taxa following the results of the phylogenomic tree resulting from the ASTRAL analysis. For this analysis, we included *C. penicillata* samples from two different biogeographic regions (the Cerrado and the ecological transition between the Atlantic Forest and Cerrado). Z scores for the ABBABABA results were calculated in R with a jackknife test using the jackKnife.R script provided with ANGSD. Absolute Z scores above 3 were used to determine significant D-statistics.

### *Callithrix* Genomic Diversity, Population Structure, and Neutrality Tests

For each species, we calculated nucleotide diversity (Pi) and Tajima’s D (Tajima, 1989) with ANGSD by using intraspecific BAM files resulting from alignment to the *Cebus imitator* genomic assembly. First, the site allele frequency (SAF) likelihood was estimated for each species where the *Cebus imitator* genomic assembly was used as both the reference and ancestral genome with the following settings: -uniqueOnly 1 -remove bads 1 - only proper pairs 1 -trim 0 -C 50 -baq 1 -P 4 -minMapQ 25 –minQ 25 -setMinDepth 1 -setMaxDepth 50 -doCounts 1 -GL 1 –doSaf 1. Then we calculated the maximum likelihood estimate of the site frequency spectrum (SFS) with the realSFS command and SAF likelihood estimates. Species per site thetas were calculated with SFS estimates set to the -pest flag, the reference and ancestral assembly set to *Cebus imitator*, and the other following parameters: uniqueOnly 1 -remove bads 1 -only proper pairs 1 - trim 0 -C 50 -baq 1 -P 4 -minMapQ 40 -minQ 32 -setMinDepth 1 -setMaxDepth 50 -doCounts 1 -GL 1 -doSaf 1 -doThetas 1. Finally, we produced sliding window estimates of Tajima’s D, nucleotide diversity, and Watternson’s theta with a window side of 50000 bp and step size of 10000 bp with the command ‘thetaStat do stat’ using the theta estimates from the previous command.

To estimate population differentiation between species, we calculated Fst estimates with ANGSD for all possible pairwise combinations of *Callithrix* species using BAMs aligned to the *Cebus imitator* genomic assembly. As a complement to Fst, we also calculated the population branch statistic (PBS) for each possible trio of *Callithrix* species using BAMs aligned to the *Cebus imitator* genomic assembly. The PBS estimates focal branch length and have been used to identify regions that have undergone a selective sweep in a focal species, and has been used in various studies as an alternative to Fst-based genome wide scans (Shpak et al., 2024). First, we used species SAF likelihood calculations described above to generate all possible species pairwise 2dsfs measures with the realSFS utility in ANGSD. Then PBS estimates were calculated for all possible three-way combinations of species (which also produces two-way Fst estimates for each species in a three-way combination) with the realSFS fst index command in combination with single species SAF likelihoods and 2dsfs estimates. We then summarized per site Fst and PBS estimates for 50000 base pair windows with 10000 base pair steps using realSFS fst stats2.

To investigate population structure between species, we conducted Principle Component Analysis (PCA) using ANGSD. Output was produced with the following parameters: -uniqueOnly 1 -remove bads 1 -only proper pairs 1 -trim 0 -C 50 -baq 1 –P 4 -minMapQ 40 -minQ 32 -setMinDepth 1 -setMaxDepth 50 - doCounts 1 -skipTriallelic 1 -doMajorMinor 1 -doMaf 1 –doIBS 1 -doCov 1 -makeMatrix 1 -GL 1 -doSaf 1. We repeated this analysis by altering levels of significance for SNPs and minor allele frequency by running all possible combinations of -SNP pval =1e-3, 1e-4,1e-6, and 1e-8 and -minMaf = 0. 1, 0.25, and 0.30. Resulting eigenvalues from PCA *.covMat files for each SNP p-value and MAF level were plotted in R.

### Heterozygosity and Runs of Heterozyosity

We used ANGSD to generate genome-wide global estimates of heterozygosity for each sampled individual after alignment to the *Cebus imitator* genomic assembly. For a global estimate, we first generated SFS estimates for each sampled individual with the following ANGSD options: -uniqueOnly 1 -remove bads 1 - only proper pairs 1 -trim 0 -C 50 -baq 1 -P 4 -minMapQ 25 -minQ 25 -dosaf 1 -gl 1. Then realSFS was used to calculate individual global heterozygosity, which was then averaged across species. For samples BJT157 and BJT169-BJT171, which had the highest genomic coverage of all sampled individuals, each sample was down-sampled to 1.5x coverage to reflect average coverage of all other sampled individuals. Down sampling was carried out with SAMTOOLS for the *Cebus imitator* assembly, and global and local heterozygosity estimates were carried out as before. The BCFTOOLs v1.8 roh command (Narasimhan et al., 2016) was used to identify runs of homozygosity (ROH) across individual WGS data with the G option set to 30, which is recommended by the software to account for genotyping error. The raw ROH results were filtered then by read group (RG) and for phred quality scores of at least 30.

### Identification, Functional Enrichment Analysis, and Functional Networks of *Callithrix* Candidate Positive Selection Genes

To look for possible targets of positive selection among *Callithrix* species, we used R to select the top scoring genetic loci in the 99.9th percentile of genome-wide windowed PBS calculations for each possible trio of *Callithrix* species. The web version of Ensembl Release 102 Biomart (Kinsella et al., 2011) was used to ‘liftover’ outlier regions mapped to *Cebus imitator* genomic assembly to the *C. jacchus* ASM275486v1 genomic assembly and identify homologous *C. jacchus* genes found within these regions. Functional and background information on identified outlier genes was obtained manually from Genecard (Stelzer et al., 2016; Safran et al., 2021) and Uniprot databases (Bateman et al., 2024). In these databases, we used human-focused information as this was the most closely related species to marmosets for which information was available. We also conducted literature searches for each gene in PUBMED to complement Genecard and Uniprot results.

Functional enrichment analysis of candidate positive selection genes for each possible PBS trio of *Callithrix* species was conducted with the g:GOSt function from the g:Profiler web-interface (Raudvere et al., 2019). Each list of candidate positive selection genes was input into the g:GOSt interface, and for options we set the organism to *C. jacchus*, which was the closest species to all of our species of interest. Under “Advanced Options,” we used Benjamini-Hochberg FDR for significance tolerance. *GMT and *GEM files were downloaded from the g:Profiler web interface and input into CYTOSCAPE 3.10.0 to make a functional network for each list of positive selection genes with the EnrichmentMap plug-in.

## Author contributions statement

JM formulated the idea for the study, collected samples, obtained funding, conducted wet and dry laboratory work, and wrote the original manuscript. VB and IOS provided field assistance and logistical support. NHAC provided field assistance and logistical support. JAD provided study guidance, and logistical support. CSI gave access and provided logistical support to collect samples from animals kept at Guarulhos Zoo. SBM provided logistical support and veterinary assistance to collect samples from animals kept at the Rio de Janeiro Primatology Center. PAN gave access and provided logistical support to collect samples from animals kept at CEMAFAUNA. LCMP gave access and provided logistical support to collect samples from animals kept at CEMAFAUNA. MP provided field assistance and logistical support. AP gave access and provided logistical support to collect samples from animals kept at CPRJ. RCHR, RAC, and CRRM were major contributors in writing the manuscript and associated with analytical methodologies. DLS assisted with associated in filed work and molecular biological work. ACS was a major contributor in writing the manuscript, provided study guidance, and logistical support. CR gave assistance in data processing, carried out divergence analysis, gave significant guidance and input in the development of this study. All authors read and approved the final manuscript.

## Acknowledgments

We would like to thank CEMAFAUNA, Rio de Janeiro Primatology Center, the Guarulhos Municipal Zoo, the Beagle Lab at UFV, the German Primate Center, and many biologists, field technicians, veterinarians, and other individuals that made this research possible. We would like to thank The Marmoset Working Group, and especially Mike Powers and Cory Ross for insightful discussion and comments on this work. We would like to especially thank Katerina Guschanski and her research group for several important discussions, insightful comments, and methodological suggestions over the course of this work. This work was supported by a Brazilian CNPq Jovens Talentos Postdoctoral Fellowship (302044/2014–0), a Brazilian CNPq DCR grant (300264/2018–6), a AAPA Professional Development Grant, Goldberg Research Grant, an American Society of Primatologists Conservation Small Grant, a Marie-Curie Individual Fellowship (AMD-793641-4), and an International Primatological Society Research Grant for JM. The funding agencies had no roles in study design, data collection and analysis, decision to publish, or preparation of the manuscript.

