## Supplementary Figures for "Genomic Insights into Derived Dwarfism and Exudivory in a Genus (*Callithrix*) of the World’s Smallest Anthropoid Monkeys"

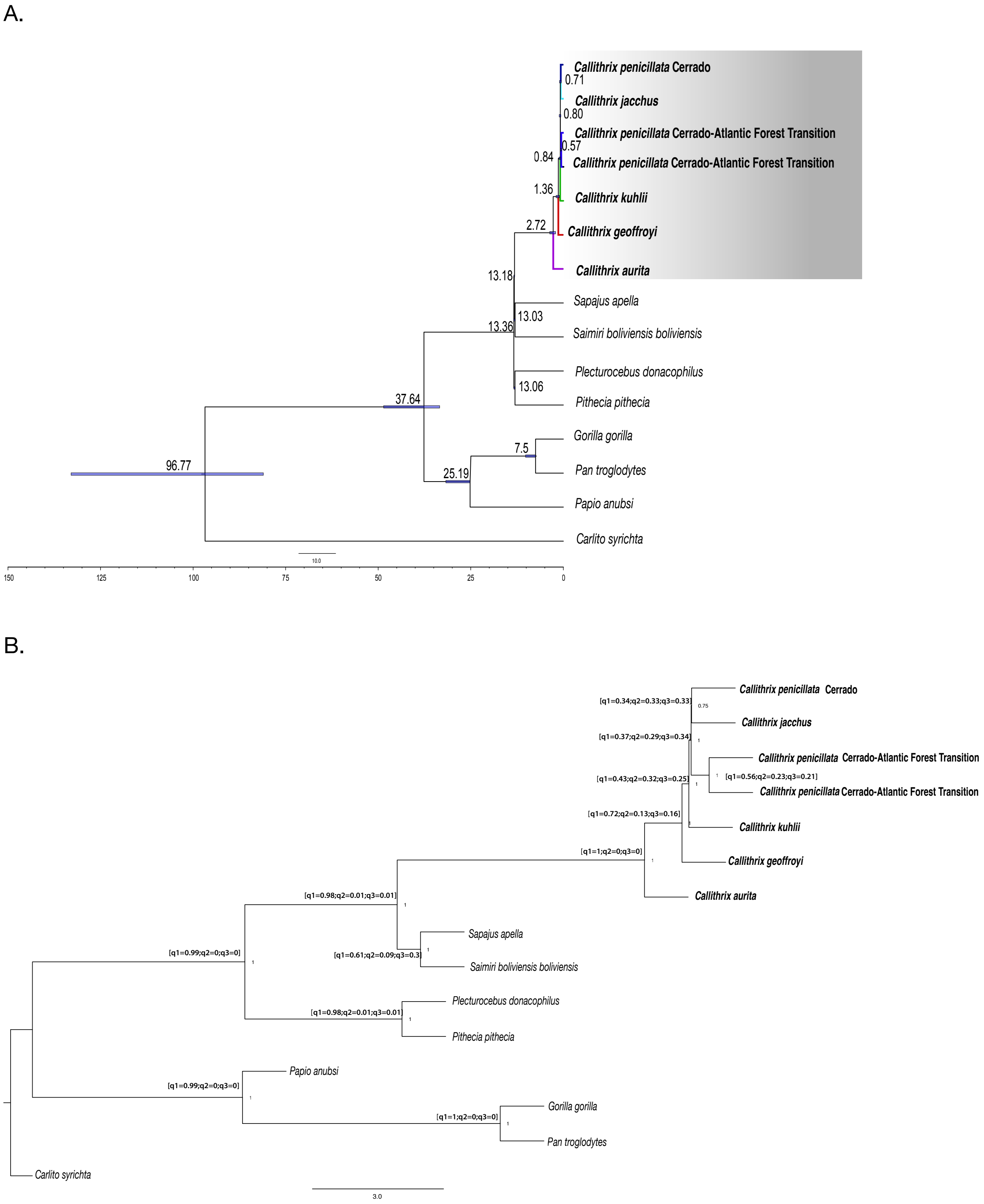


Supplementary Figure S1. *Callithrix* phylogenetic trees. A. *Callithrix* multispecies coalescent-based ASTRAL species tree and divergence time estimates as millions of years ago (MYA) shown at major nodes. Blue bars at nodes represent 95% divergence time confidence intervals. B. *Callithrix* multispecies coalescent-based ASTRAL species tree. ASTRAL quartet scores are shown on the left side of each node. Branch Support in Local Posterior Probability (localPP) is shown on the right side of each node.

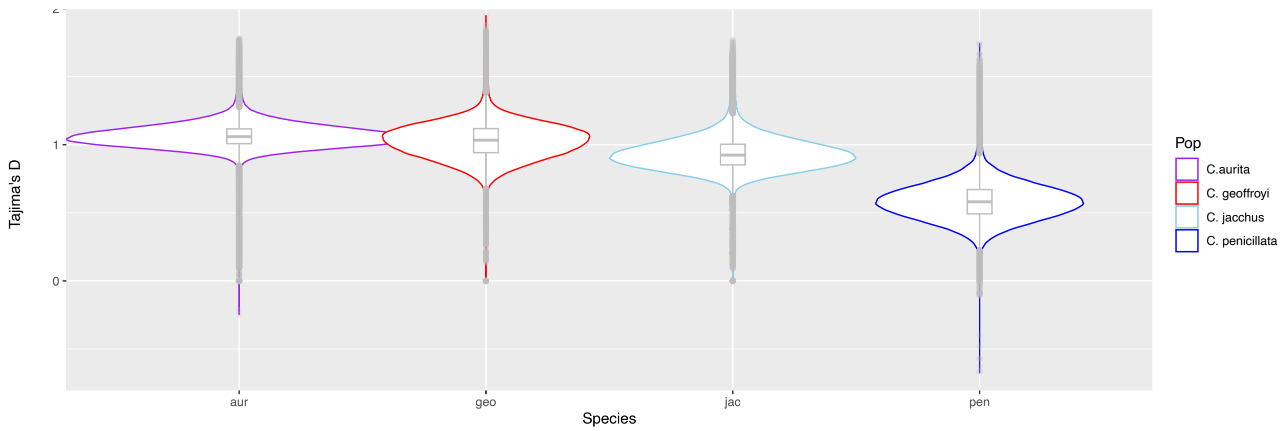


Supplementary Figure S2. Violin plots of average genome-wide Tajimas' D values for individual *Callithrix* species. The bottom box lines represent 25th percentiles, the mid-lines of boxes represent 50th percentiles/medians, and top box lines represent 75th percentiles. Figure legend of "Pop", which is an abbreviation for population, seen on the right side shows individual colors representing each species. Abbreviations along the x-axis correspond to species names as follows: aur- *C. aurita*, geo- *C. geoffroyi*, jac- *C. jacchus*, pen- *C. penicillata*

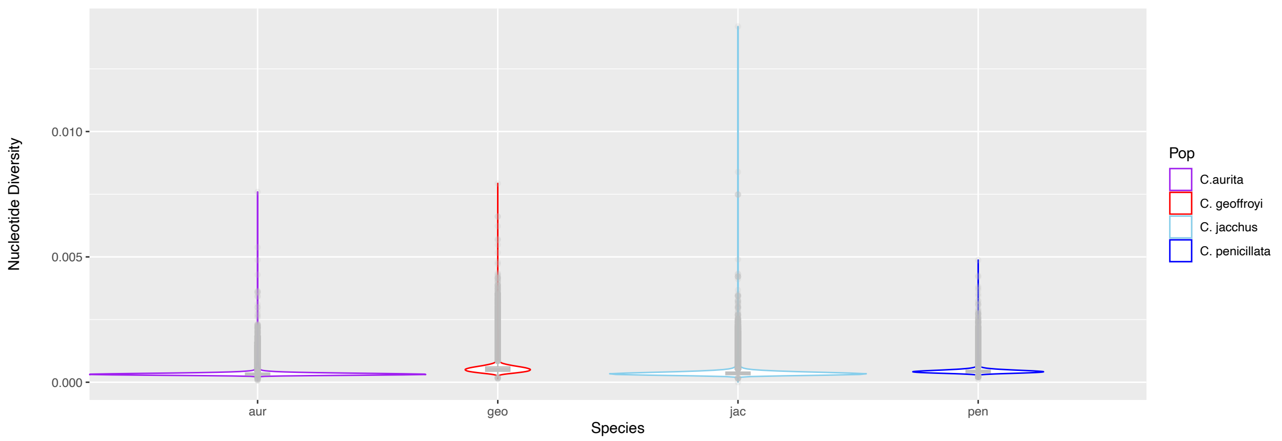


Supplementary Figure S3. Violin plots of average genome-wide nucleotide diversity values for individual *Callithrix* species. The bottom box lines represent 25th percentiles, the mid-lines of boxes represent 50th percentiles/medians, and top box lines represent 75th percentiles. Figure legend of "Pop", which is an abbreviation for population, seen on the right side shows individual colors representing each species. Abbreviations along the x-axis correspond to species names as follows: aur- *C. aurita*, geo- *C. geoffroyi*, jac- *C. jacchus*, pen- *C. penicillata.*

A

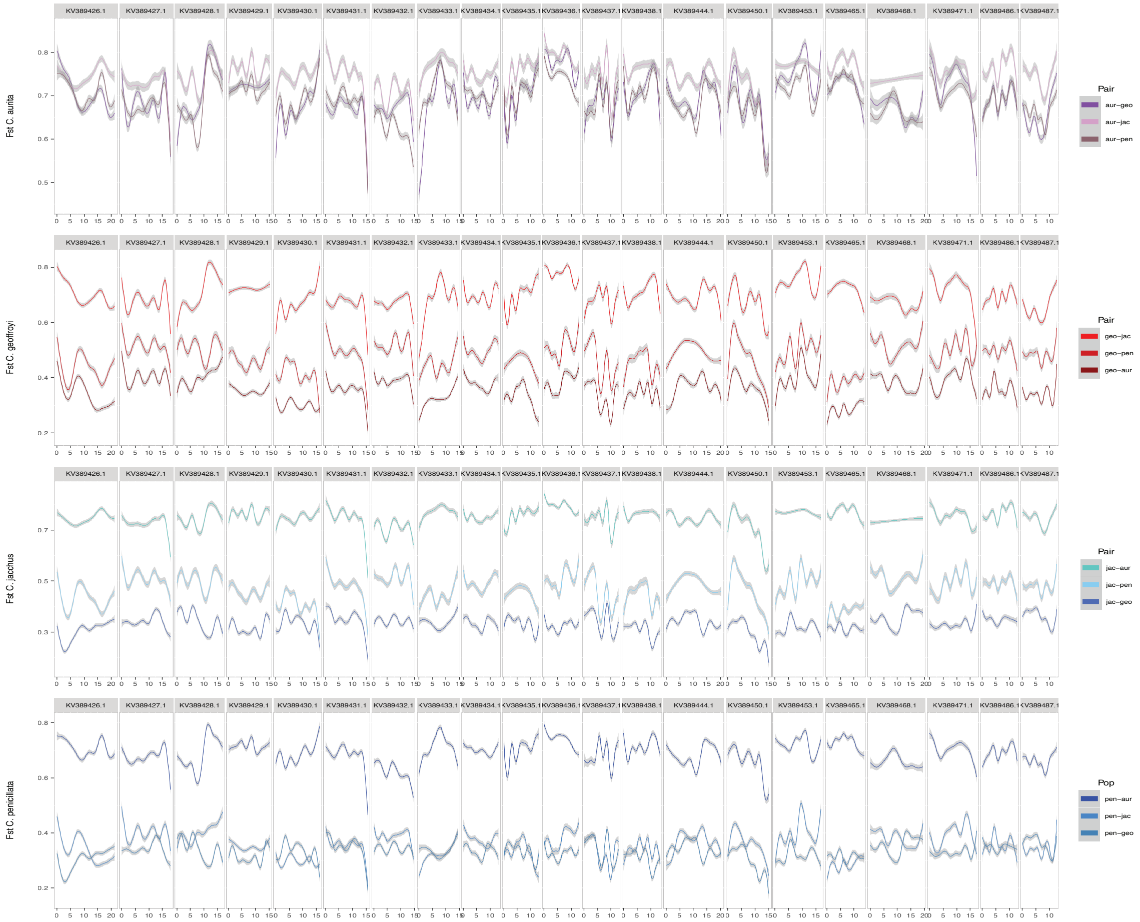


Supplementary Figure S4. Pair-wise comparisons of Fst values for genome-wide genomic windows between all possible pairs of *Callithrix* species. Each plot panel features comparisons between a "focal" species and the remaining other species. Smoothed Fst values from genomic windows are shown along the 21 largest scaffolds in the *Cebus imitator* genome. Abbreviations in each legend called "Pop" (which is short for population) correspond to species names as follows: aur- *C. aurita*, geo- *C. geoffroyi*, jac- *C. jacchus*, pen- *C. penicillata.*

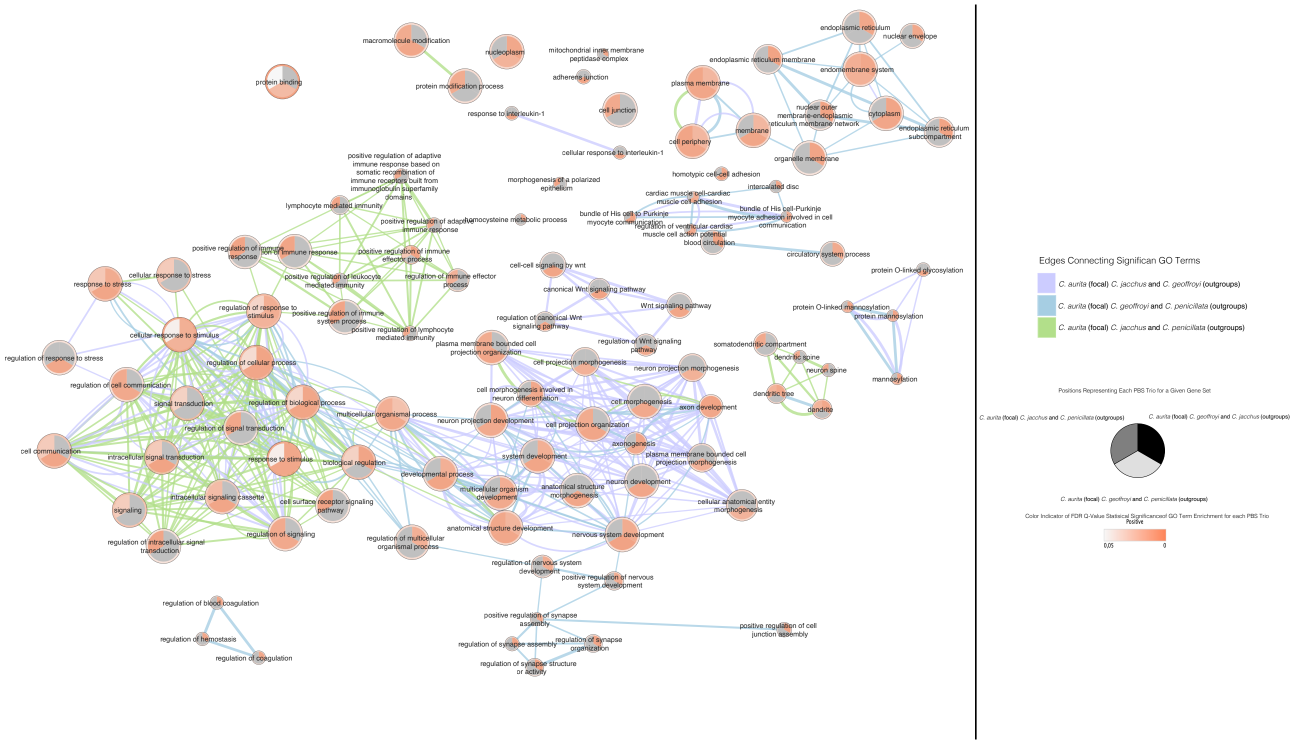


Supplementary Figure S5. Cytoscape Enrichment Map of significantly enriched  GO terms from all PBS trios in which *C. aurita* is the focal species. Each circle represents a set of genes for a given GO term, and each trio is represented by a given position in each circle. *Callithrix aurita/C. jacchus/C. geoffroyi* are shown in the upper right, *C. aurita/C. jacchus/C. penicillata* are shown in the upper left of the gene set circle, and *C. aurita/C. geoffroyi/C. penicillata* are shown in the bottom third of the gene set circle. A grey color within each gene set circle signifies lack of significant enrichment for a given GO term. Otherwise, the statistical significance of the GO term enrichment is signified by increasing intensity of red, which signifies a range of FDR Q-values from 0.05 to 0. Edge lines show connections between GO terms and are color-coded for each PBS trio, with *C. aurita/C. geoffroyi/C. penicillata* represented by light blue, *C. aurita/C. jacchus/C. penicillata* by light green, and *C. aurita/C. jacchus/C. geoffroyi* by light purple.

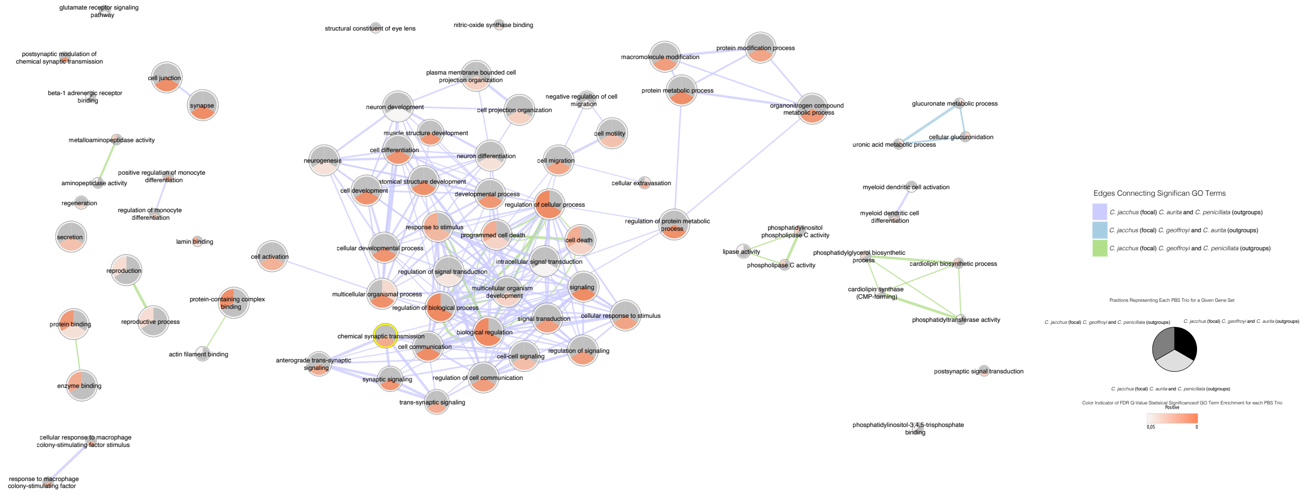


Supplementary Figure S6. Cytoscape Enrichment Map of significantly enriched GO terms from all PBS trios in which *C. jacchus* is the focal species. Each circle represents a set of genes for a given GO term, and each trio is represented by a given position in each circle. *Callithrix jacchus/C. aurita/C. geoffroyi* are shown in the upper right, *C. jacchus/C. geoffroyi/C. penicillata* are shown in the upper left of the gene set circle, and *C. jacchus/C. aurita/C. penicillata* are shown in the bottom third of the gene set circle. A grey color within each gene set circle signifies lack of significant enrichment for a given GO term. Otherwise, the statistical significance of the GO term enrichment is signified by increasing intensity of red, which signifies a range of FDR Q-values from 0.05 to 0. Edge lines show connections between GO terms and are color-coded for each PBS trio, with *C. jacchus/C. geoffroyi/C. penicillata* represented by light green, *C. jacchus/C. aurita/C. penicillata* by light purple, and *C. jacchus/C. aurita/C. geoffroyi* by light blue.

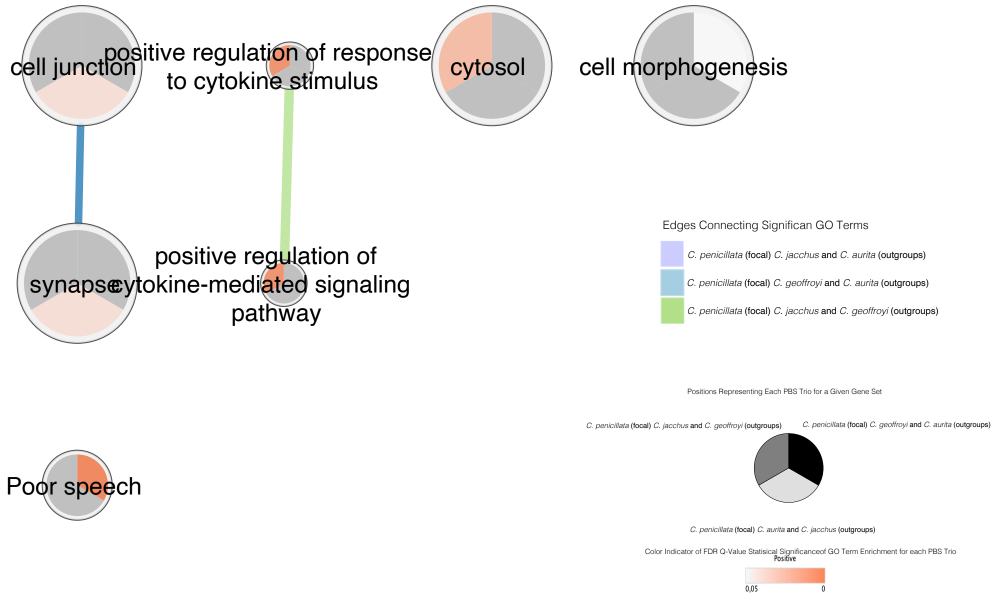


Supplementary Figure S7. Cytoscape Enrichment Map of significantly enriched GO terms from all PBS trios in which *C. penicillata* is the focal species. Each circle represents a set of genes for a given GO term, and each trio is represented by a given position in each circle. *Callithrix penicillata/C. aurita/C. geoffroyi* are shown in the upper right, *C. penicillata/C. geoffroyi/C. jacchus* are shown in the upper left of the gene set circle, and *C. penicillata/C. aurita/C. jacchus* are shown in the bottom third of the gene set circle. A grey color within each gene set circle signifies lack of significant enrichment for a given GO term. Otherwise, the statistical significance of the GO term enrichment is signified by increasing intensity of red, which signifies a range of FDR Q-values from 0.05 to 0. Edge lines show connections between GO terms and are color-coded for each PBS trio, with *C. penicillata/C. geoffroyi/C. jacchus* represented by light green, *C. penicillata/C. aurita/C. jacchus* by light purple, and *C. penicillata/C. aurita/C. geoffroyi* by light blue.
